# Alcohol-Evoked Calcium signalling Drives Distinct Responses in Zebrafish Hepatocytes and Pancreatic Acinar Cells

**DOI:** 10.64898/2026.08.21.742910

**Authors:** Kaushani Ghosh, Macarena Pozo-Morales, Sema Elif Eski, Abhishek Tanwar, Rajender K Motiani, Sumeet Pal Singh

**Author notes:** These authors contribute equally to the work (Kaushani Ghosh, Macarena Pozo-Morales).

## Abstract

Alcohol exposure perturbs intracellular calcium (Ca²⁺) homeostasis in digestive organs, yet whether common or organ-specific mechanisms coordinate this response remains unclear. Using an acute ethanol paradigm in zebrafish, single-cell transcriptomics revealed broad upregulation of Ca²⁺-signalling genes in hepatocytes and pancreatic acinar cells. In vivo Ca²⁺ buffering with SpiCee, a genetically encoded chelator, demonstrated a shared requirement for Ca²⁺ flux: in hepatocytes, lineage-restricted buffering was associated with pronounced cytoplasmic vacuolation composed of lipid-negative vesicles, consistent with stalled lysosomes or autophagosomes; in pancreatic acinar cells, it was associated with accumulation of aggregated/misfolded protein. Mechanistic experiments using pharmacological inhibitors implicated distinct molecular contributors in each tissue. In hepatocytes, inhibition of Pikfyve or its downstream effector, the lysosomal Ca²⁺ channel TRPML1, phenocopied Ca²⁺ buffering. While, in acinar cells, Pick1 inhibition produced analogous associations. These data position Pikfyve and Pick1 as organ-specific components linked to the Ca²⁺-coupled alcohol response. Notably, pharmacologic activation of TRPML1 in hepatocytes recapitulated alcohol-like Ca²⁺ dynamics but increased macrophage recruitment and cell death, indicating that Ca²⁺ signalling is required for the alcohol response yet can be detrimental when amplified. Together, our results support a model in which alcohol elicits a shared Ca²⁺ dynamics across liver and pancreas, modulated by tissue-specific molecular nodes.

**Graphical Abstract:** 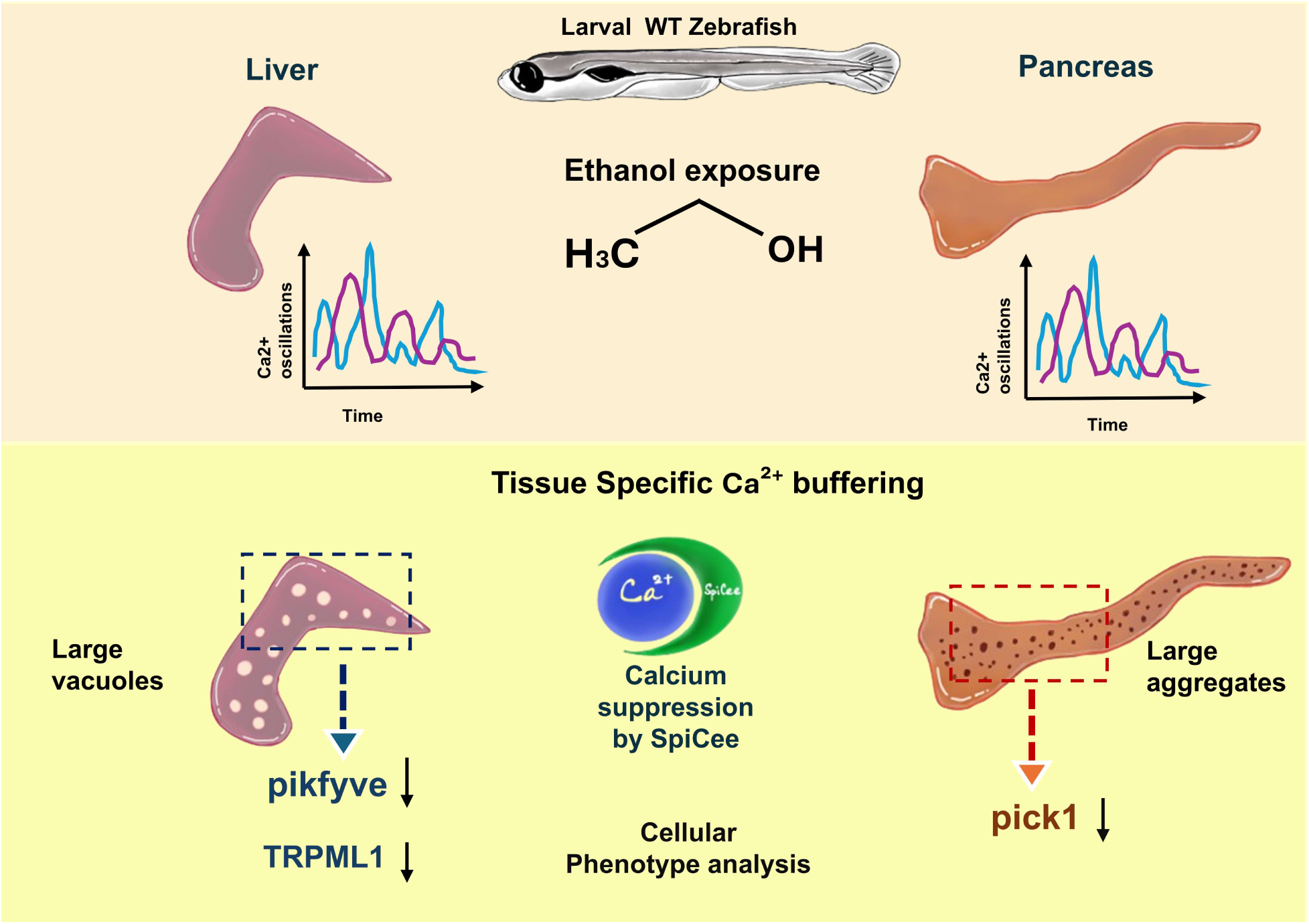

## Introduction

Alcohol consumption is a major global health concern ^1,2^, with the liver and exocrine pancreas among its principal targets. Chronic exposure can cause alcohol-associated liver disease (ALD), ranging from steatosis to hepatitis, fibrosis, and cirrhosis, as well as acute and chronic pancreatitis. These pathologies frequently coexist, and disease in one organ is associated with an increased likelihood of injury in the other ^3–5^. However, the early cellular mechanisms through which alcohol affects these organs, and whether they are shared or tissue-specific, remain incompletely understood.

In hepatocytes, ethanol metabolism generates acetaldehyde and reactive oxygen species, promoting oxidative stress, mitochondrial and endoplasmic reticulum dysfunction, impaired protein homeostasis, and cell death ^6^. Repeated injury activates hepatic stellate cells and drives fibrosis ^7^. In pancreatic acinar cells, alcohol and its metabolites promote pathological calcium signals, premature activation of digestive zymogens, cellular autodigestion, and inflammation ^8^. Recurrent injury can progress to chronic pancreatitis, with fibrosis and loss of acinar tissue ^9^.

Disruption of intracellular calcium homeostasis is an early feature of alcohol-induced injury in both organs. In hepatocytes, ethanol alters calcium signalling and communication between intracellular stores, affecting metabolism, organelle function, and stress responses ^10,11^. In pancreatic acinar cells, ethanol induces abnormal global cytosolic calcium signals that impair ATP production, vesicular trafficking, and zymogen control ^12^. Thus, calcium dysregulation may represent a common upstream response to alcohol exposure.

A major unresolved question is whether this shared calcium response produces similar cellular consequences in hepatocytes and acinar cells or is interpreted through distinct tissue-specific pathways. It is also unclear whether alcohol-evoked calcium signalling is protective, harmful, or dependent on signal magnitude and cellular context. Moreover, the molecular pathways linking altered calcium dynamics to organ-specific pathology remain poorly defined.

Here, we used larval zebrafish to compare calcium-dependent responses to acute ethanol exposure in hepatocytes and pancreatic acinar cells. By combining single-cell RNA sequencing, tissue-specific calcium reporters, live imaging, genetically encoded calcium buffering, and pharmacological perturbation, we examined both the shared requirement for calcium signalling and the mechanisms underlying distinct cellular outcomes. Our analyses identify the PIKfyve– TRPML1 endolysosomal pathway in hepatocytes and PICK1-associated vesicular regulation in acinar cells as tissue-specific components of the alcohol response.

## Results

### Single-cell transcriptomics identifies ethanol-responsive calcium signalling programs in hepatocytes and pancreatic acinar cells

To define the cellular landscape of the larval zebrafish digestive system and assess its response to alcohol exposure, we performed single-cell RNA sequencing using the 10x Genomics platform on cells dissociated from the dissected gut region of control 6 days post-fertilization (dpf) larvae and larvae treated with 2% ethanol for 16 hours (**Figure 1A**). Integrated UMAP analysis resolved major organ-associated populations corresponding to liver, pancreas, intestine, kidney, proliferating cells, and cell types shared across multiple organs (**Figure 1B**).

**Figure 1:**
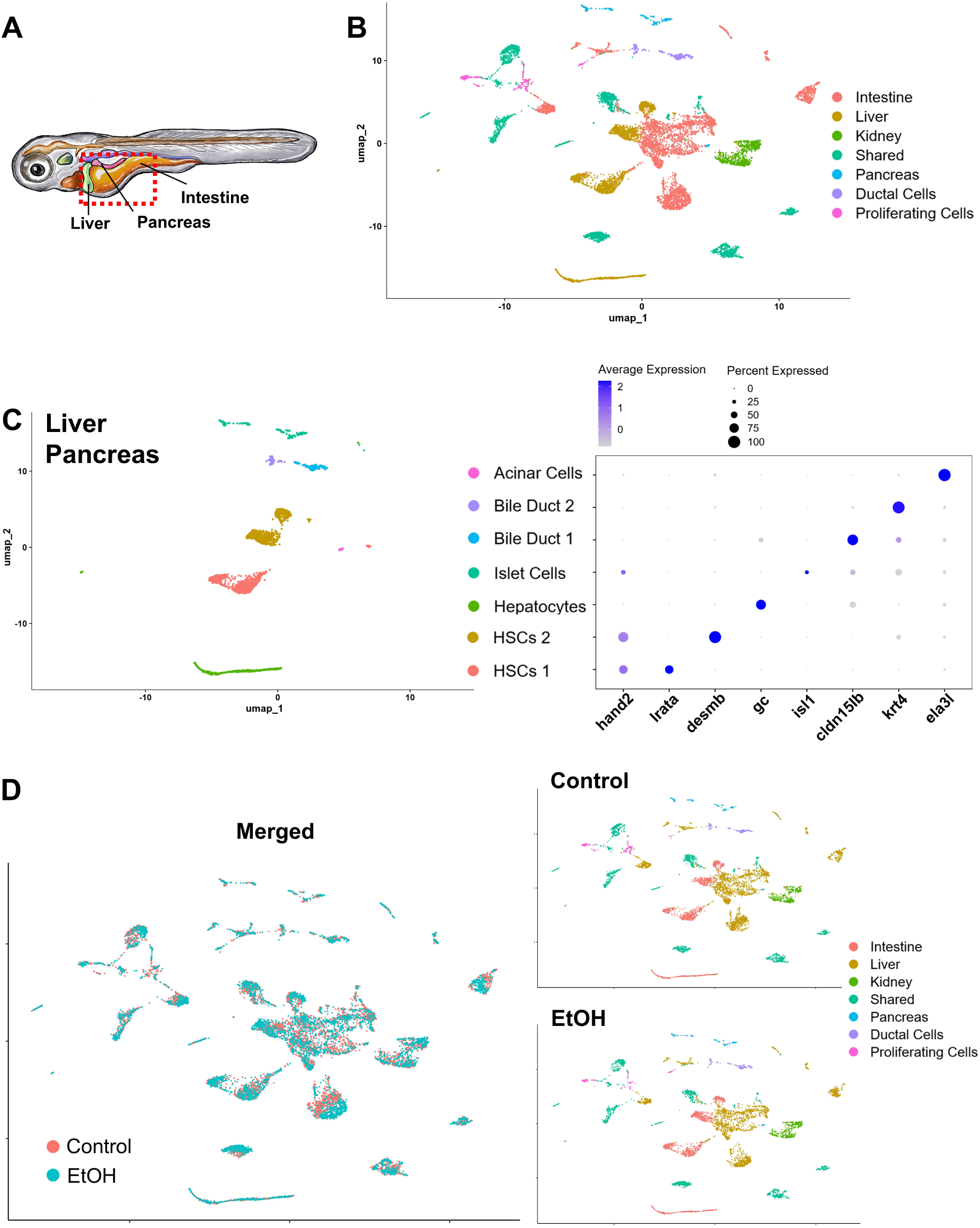
Single-cell transcriptomic atlas of zebrafish digestive organs. **(A)** Schematic representation of the zebrafish larva indicating the anatomical region encompassing the liver, pancreas, intestine, and adjacent tissues that were dissected for single-cell RNA sequencing analysis. The dashed red box highlights the region collected for sequencing. **(B)** Single cell atlas of Zebrafish showing major cellular populations identified across the dissected digestive organs. Cells were clustered and annotated based on established marker gene expression, revealing organ-specific populations corresponding to the intestine, liver, kidney, pancreas, ductal cells, proliferating cells, and immune cell populations. **(C)** Sub-clustering analysis of liver and pancreas-derived cells. Dot plots display representative marker genes used for cluster annotation, where color intensity indicates average gene expression and dot size represents the percentage of cells expressing each marker. **(D)** UMAP visualization of the integrated single-cell RNA-sequencing dataset showing cells from control and 2% ethanol-treated larvae. Left, merged UMAP with cells colored according to treatment condition. Middle and right, the same UMAP embedding shown separately for control and ethanol-treated cells, respectively.

Within the liver–pancreas compartment we identified seven transcriptionally distinct sub-populations, including hepatocytes, pancreatic acinar cells, islet cells, two bile duct populations, and two hepatic stellate cell populations (**Figure 1C**). These identities were supported by canonical marker expression: hepatocytes expressed *gc*, acinar cells expressed *ela3l*, bile duct cells expressed *cldn15lb* and *krt4*, and islet cells expressed *isl1*. The two hepatic stellate cell populations were distinguished by differential expression of *hand2*, *lrata*, and *desmb*.

Sub-clustering of the intestine–kidney compartment resolved eight populations, including enterocytes, intestinal stem cells, tuft cells, enteroendocrine cells, Best4+ cells, podocytes, and two stromal populations (**Supplementary Figure 1A**). Finally, sub-clustering of the shared cell compartment identified vascular endothelial cells, lymphatic endothelial cells, pericytes, erythrocytes, macrophages/immune cells, lymphocytes, proliferating cells, and fibroblasts (**Supplementary Figure 1B**). These populations were annotated using established lineage markers, including *mpeg1.1* for macrophages, *fli1* and *abcc9* for vascular endothelial cells, *klf1* for erythrocytes, *il4* for lymphocytes, *mki67* for proliferating cells, *lyve1b* for lymphatic endothelial cells, and *prrx1a/prrx1b* for fibroblasts. Notably, the proliferating-cell cluster showed enrichment of the macrophage marker mpeg1.1, indicating that a substantial proportion of proliferating cells were macrophage-lineage cells (**Supplementary Figure 1C, D**). Together, this atlas provided a cell-type-resolved framework to examine how acute alcohol exposure affects digestive organs.

To assess whether ethanol exposure altered the overall representation of cellular populations within the dataset, we visualized control and ethanol-treated cells together and separately on the integrated UMAP embedding (**Figure 1D**). Cells from both conditions were broadly distributed across the major populations, with substantial overlap between control and ethanol-treated cells in the merged representation, suggesting that no unique cell population was present in either condition.

We next investigated the transcriptional response to acute ethanol exposure by comparing larvae treated with 2% ethanol for 16 hours to untreated controls across all annotated cell populations. Differential gene expression analysis for the entire dataset revealed 252 genes upregulated and 154 genes downregulated in the cells from ethanol-treated larvae (**Figure 2A, Supplementary Table 1**). Among the most strongly induced genes, *fkbp5* was notable because it was the only gene consistently upregulated across all annotated cell types examined (**Figure 2B**). This broad induction identifies *fkbp5* as a pan-cellular marker of the acute ethanol response in the larval digestive system. Importantly, *fkbp5* encodes FKBP51, a stress-responsive immunophilin that has been linked to Ca²⁺ homeostasis; previous studies have shown that FKBP51 can regulate store-operated Ca²⁺ entry and modulate Ca²⁺ entry-induced cellular phenotypes ^13,14^.

**Figure 2:**
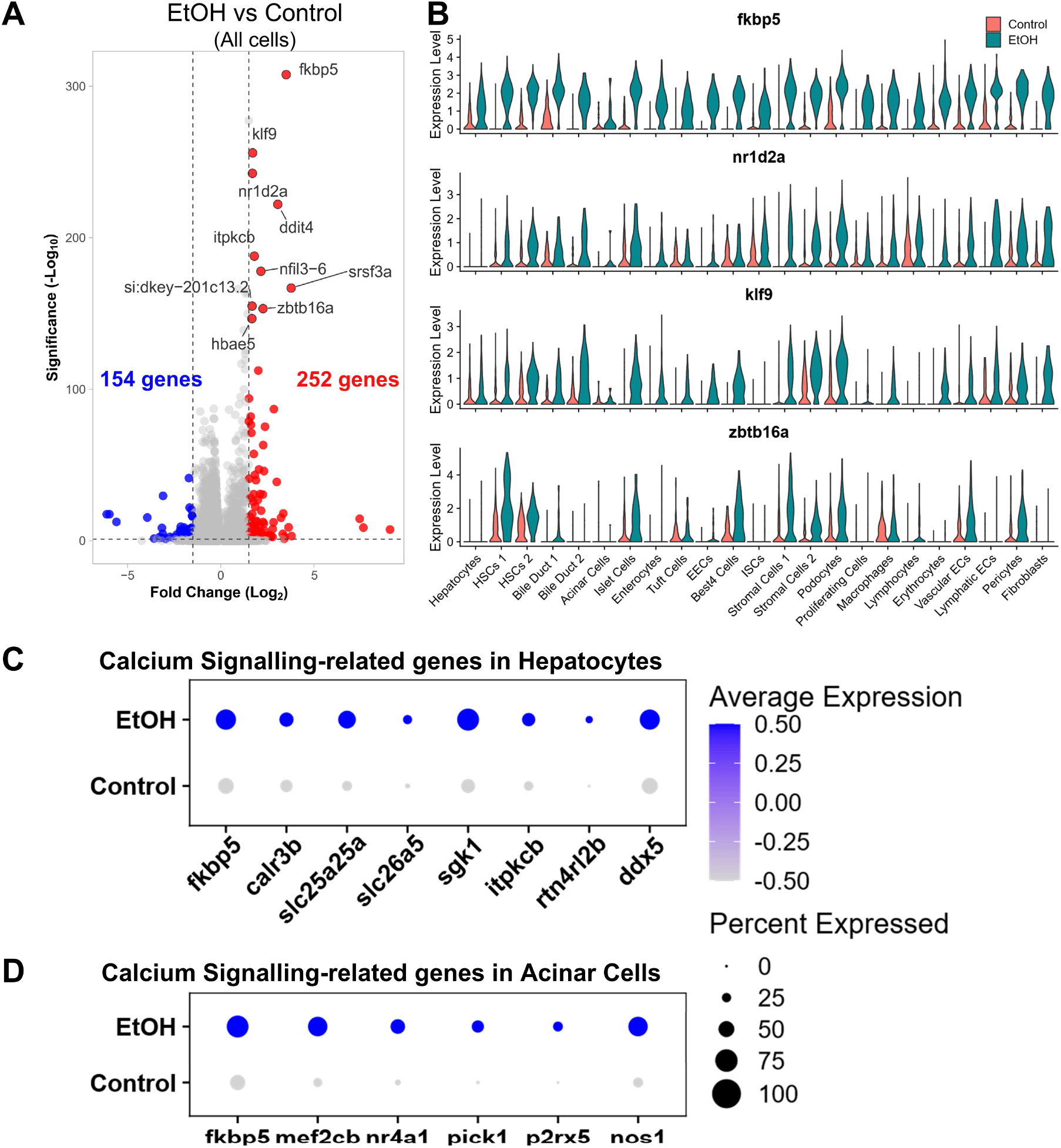
Ethanol exposure induces stress-responsive transcriptional programs and calcium signalling-associated gene expression in hepatocytes and pancreatic acinar cells. **(A)** Volcano plot showing differential gene expression between ethanol-treated (EtOH) and control cells from the single-cell RNA sequencing dataset. Significantly upregulated genes in ethanol-treated samples are shown in red, while downregulated genes are shown in blue. The x-axis represents log2 fold change, and the y-axis represents statistical significance (-log₁₀ adjusted *P* value). **(B)** Violin plots depicting the expression of representative ethanol-responsive genes (*fkbp5, nr1d2a, klf9,* and *zbtb16a*) across annotated cell populations with the comparison of Control and Ethanol treated samples. **(C)** Dot plot shows the expression of calcium signalling-associated genes in hepatocytes from control and ethanol-treated larvae. Ethanol exposure increased the expression of several calcium-responsive genes, including fkbp5, calr3b, slc25a25a, slc26a3, sgk1, itpkcb, rtn4rl2b, and ddx5. Dot size indicates the percentage of expressing cells, while color intensity represents average gene expression. **(D)** Dot plot showing calcium signalling-associated gene expression in pancreatic acinar cells following ethanol exposure. Ethanol-treated acinar cells exhibited elevated expression of genes implicated in calcium homeostasis and signalling, including *fkbp5, mef2cb, nr4a1, pick1, p2rx5,* and *nos1*.

Next, we focused on hepatocytes and pancreatic acinar cells, two digestive cell types central to alcohol-associated liver disease and pancreatitis. In hepatocytes, ethanol exposure induced a stress-response signature in hepatocytes (**Supplementary Figure 2**), consistent with previous findings in zebrafish and mice ^15–17^. Further, we also observed an enrichment for cellular calcium ion homeostasis by Gene Ontology (GO) analysis (**Supplementary Figure 2A**), which included the significant increase in the expression of multiple Ca²⁺ signalling-associated genes, including *fkbp5*, *calr3b*, *slc25a25a*, *slc26a5*, *sgk1*, *itpkcb*, *rtn4rl2b*, and *ddx5* (**Figure 2C**). These genes are associated with diverse aspects of Ca²⁺ homeostasis, including endoplasmic reticulum Ca²⁺ buffering, mitochondrial Ca²⁺ transport, ion regulation, and inositol phosphate-mediated Ca²⁺ signalling^13,18–20^.

A parallel response was observed in pancreatic acinar cells. Ethanol-treated acinar cells showed increased expression of Ca²⁺ signalling-associated genes, including *fkbp5*, *mef2cb*, *nr4a1*, *pick1*, *p2rx5*, and *nos1* (**Figure 2D**). These findings indicate that acute ethanol exposure induces a broad systemic stress program across digestive and associated tissues, while converging on Ca²⁺ signalling-associated transcriptional responses in hepatocytes and pancreatic acinar cells. This cell-type-resolved response provides the basis for testing how Ca²⁺ signalling contributes to alcohol-induced phenotypes in the liver and pancreas.

### Ethanol induces asynchronous calcium oscillations in pancreatic acinar cells of zebrafish larva

To test whether the ethanol-induced calcium-associated transcriptional program observed in pancreatic acinar cells corresponded to altered calcium activity *in vivo*, we generated a new transgenic reporter line expressing the genetically encoded calcium indicator GCaMP6s in acinar cells, *Tg(ela3l:GCaMP6s)*.

Using this line, we performed live time-lapse confocal imaging to directly visualize calcium dynamics in the exocrine pancreas following ethanol exposure (**Figure 3A**). Imaging was performed for 12 minutes. In control, 6 dpf larvae, acinar cells showed limited changes in GCaMP6s fluorescence over the imaging period (**Supplementary Video 1**). Similar low-baseline activity was observed after feeding (data not shown). In contrast, 6 dpf larvae exposed to 2% ethanol for 16 - 18 hours displayed dynamic changes in GCaMP6s fluorescence within the acinar cell population (**Supplementary Video 2**). Quantification showed that ethanol exposure significantly increased the number of oscillating acinar cells per animal compared with controls (p-value = 0.010, Mann–Whitney U test) (**Figure 3B**). These fluorescence changes appeared as transient calcium signals that occurred asynchronously among individual acinar cells, rather than as a coordinated response across the entire tissue (**Figure 3C**).

**Figure 3:**
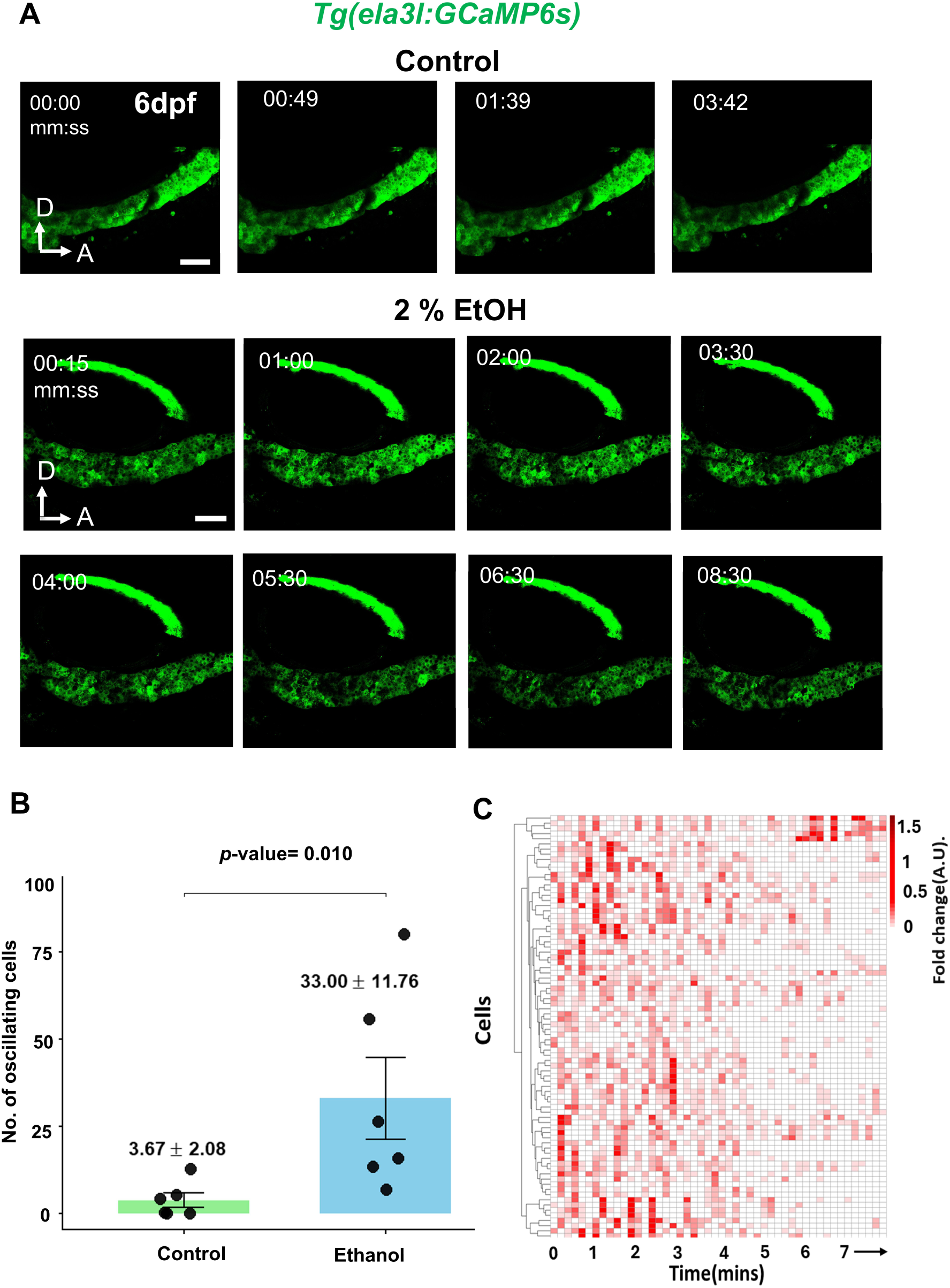
Ethanol induces asynchronous calcium oscillations in pancreatic acinar cells. **(A)** Time-lapse confocal images of pancreatic acinar cells from 6 days post fertilization (dpf) *Tg(ela3l:GCaMP6s)* zebrafish larvae under control conditions. Time stamps indicate minutes:seconds (mm:ss). Scale bar 50μm. Time-lapse images of pancreatic acinar cells following exposure to 2% ethanol (EtOH) for 16 - 18 hours. Scale bar 50μm. **(B)** Quantification of the number of oscillating pancreatic cells in eight minutes of imaging in control and ethanol-treated larvae. Data are presented as Mean ± SEM. Each dot represents one biological replicate. Statistical analysis was performed using an unpaired two-tailed Mann– Whitney U test (Wilcoxon rank-sum test). p-value = 0.010. **(C)** Heatmap representation of calcium dynamics in individual pancreatic acinar cells over time following ethanol treatment. Each row represents a single acinar cell and every column represents a single frame of time lapse imaging. Color intensity corresponds to normalized fluorescence fold change (ΔF/F or arbitrary units, A.U.), indicating calcium activity.

This pattern is consistent with ethanol-induced spontaneous calcium activity previously observed in zebrafish hepatocytes ^11^. Further, similar to the inflammatory response previously observed in the zebrafish liver following alcohol exposure, ethanol exposure also induced macrophage infiltration to the exocrine pancreas (**Supplementary Figure 3**) ^21^. Together with the hepatocyte response, these findings indicate that acute ethanol exposure induces calcium-associated transcriptional and functional responses in both the liver and exocrine pancreas.

### SpiCee-mediated calcium buffering enhances ethanol-induced hepatocyte vacuolation

The transcriptional dysregulation of calcium-signalling pathways following ethanol exposure prompted us to investigate the functional role of intracellular calcium dynamics in hepatocytes *in vivo*. We used 6 dpf *Tg(fabp10a:EGFP)* zebrafish larvae, in which hepatocytes are specifically labelled with GFP, together with Nile Red staining to visualize neutral lipid-containing compartments. Control larvae displayed normal hepatocyte morphology with scattered Nile Red-positive lipid droplets. Following exposure to 2% ethanol, 3 of 16 larvae developed intracellular vacuole-like structures within hepatocytes (**Figure 4A**). These structures were largely negative for Nile Red, indicating that they were distinct from neutral lipid droplets.

**Figure 4:**
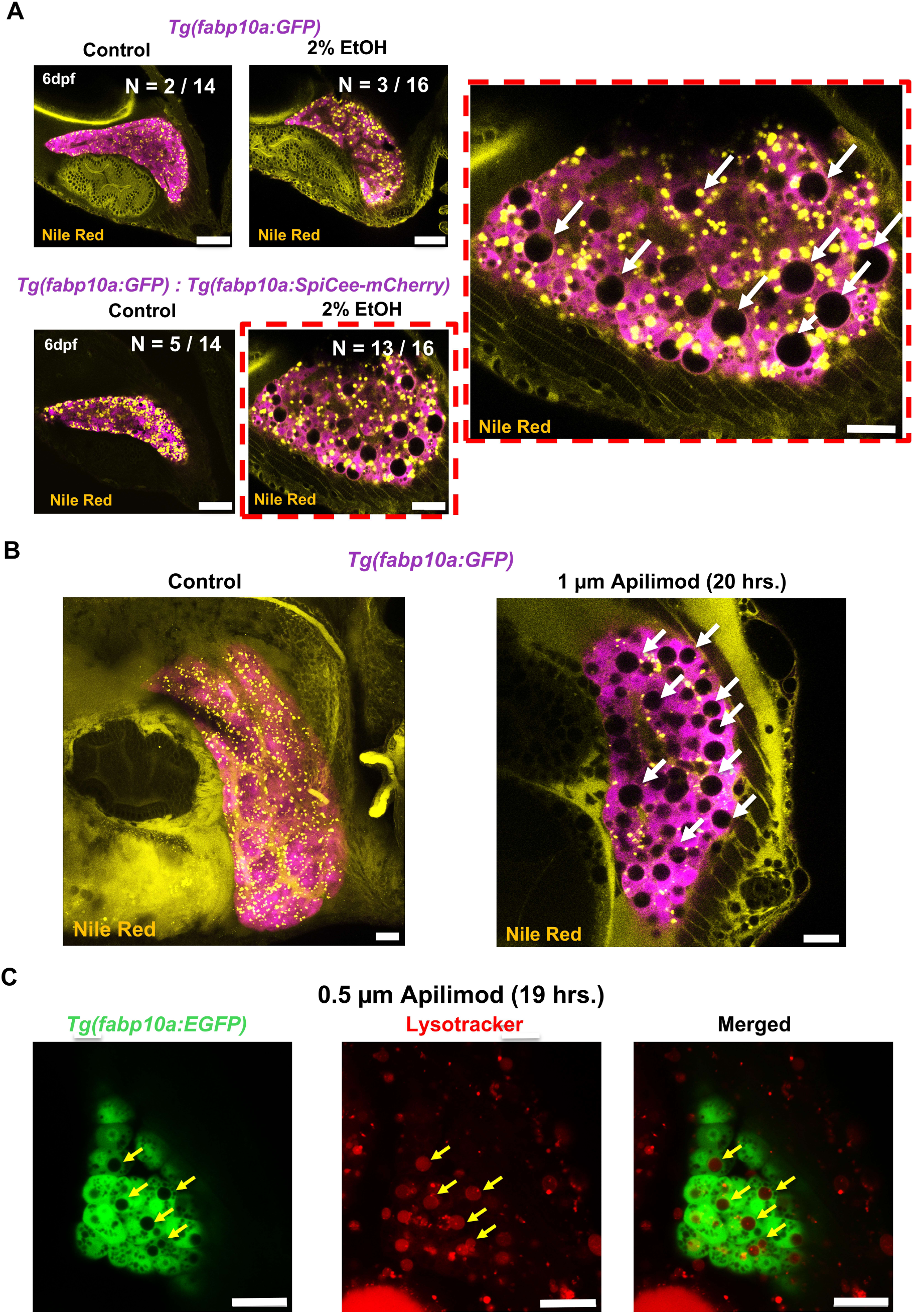
SpiCee-mediated calcium chelation induces a vacuolar liver phenotype associated with lysosomal dysfunction. **(A)** Representative confocal images of livers from 6 dpf *Tg(fabp10a:GFP)* zebrafish larvae stained with Nile Red. Control larvae displayed normal hepatic morphology, whereas exposure to 2 % ethanol (EtOH) induced the formation of numerous large intracellular vacuolar structures (white arrows) within hepatocytes. Co-expression of the calcium-chelating construct *Tg(fabp10a:SpiCee-mCherry)* markedly enhanced ethanol-induced vacuolar formation, resulting in a severe vacuolated phenotype. The dashed red box indicates the region shown at higher magnification. The incidence of the vacuolar phenotype is indicated as numbers for each image where the cutoff values for these vacuoles are set at 10. Nile Red staining labels neutral lipid-containing structures. Scale bar 50um for all panels. **(B)** Pharmacological inhibition of PIKfyve signalling using Apilimod phenocopied the ethanol-induced hepatic vacuolar phenotype. Representative confocal images of livers from *Tg(fabp10a:GFP)* larvae treated with 1 μM Apilimod for 20 h revealed the appearance of enlarged intracellular vacuoles (white arrows) compared with untreated controls. Scale bar 50um for all panels. **(C)** Lysosomal characterization of Apilimod-induced vacuoles. Representative confocal images of *Tg(fabp10a:EGFP)* larvae treated with 0.5 μM Apilimod for 19 h and stained with LysoTracker Red. Arrows indicate enlarged vacuolar structures that exhibit LysoTracker accumulation, demonstrating lysosomal association of the vacuoles. The merged image shows colocalization between hepatocyte GFP signal and LysoTracker-positive compartments. Scale bars, 20 μm.

To determine whether altered calcium signalling contributed to this phenotype, we examined *Tg(fabp10a:EGFP)*; Tg(*fabp10a:SpiCee-mCherry*) larvae expressing the genetically encoded calcium buffer SpiCee specifically in hepatocytes ^11,22,23^. SpiCee-expressing larvae showed some baseline hepatocyte vacuolation in the absence of ethanol, with vacuoles observed in 5 of 14 larvae. In contrast, ethanol exposure increased the frequency and severity of vacuolation, with numerous enlarged intracellular vacuoles observed in 13 of 16 SpiCee-expressing larvae (**Figure 4A**). Thus, hepatocyte-specific calcium buffering sensitized hepatocytes to ethanol-induced vacuolation (p-value = 0.0029, Fisher’s Exact Test). These findings indicate that intracellular calcium signalling is required to maintain hepatocyte homeostasis during ethanol exposure and suggest that disruption of this response compromises intracellular vesicle or organelle processing.

### PIKfyve inhibition induces hepatocyte vacuolation associated with lysosomal compartments

The vacuolar phenotype observed following ethanol exposure and calcium buffering resembled the enlarged cytoplasmic vacuoles previously reported after inhibition of PIKfyve ^24,25^. Consistent with this, loss-of-function mutations in zebrafish *fig4a*, which encodes a component of the PIKfyve complex, cause robust hepatic vacuolation associated with abnormal lysosomal storage and accumulation of autophagic intermediates ^26^. PIKfyve is a lipid kinase that generates phosphatidylinositol 3,5-bisphosphate [PI(3,5)P₂], an important regulator of endolysosomal membrane dynamics and vesicular trafficking ^27,28^. To determine whether disruption of this pathway could reproduce the hepatocyte phenotype, we treated 4 dpf *Tg(fabp10a:EGFP)* larvae with 1 μM apilimod, a well-established PIKfyve inhibitor, for 20 hours. Vehicle-treated larvae displayed normal hepatocyte morphology, whereas apilimod-treated larvae developed numerous enlarged intracellular vacuoles (**Figure 4B**). Thus, PIKfyve inhibition phenocopied the vacuolation observed following ethanol exposure and hepatocyte-specific calcium buffering, implicating defective endolysosomal trafficking in this phenotype.

To examine the relationship between these vacuoles and lysosomal compartments, *Tg(fabp10a:EGFP)* larvae were treated with 0.5 μM apilimod for 19 hours and stained with LysoTracker Red, which accumulates in acidic organelles ^29^. LysoTracker-positive puncta were closely associated with the enlarged vacuoles and were frequently observed along their boundaries (**Figure 4B**). These findings indicate that the apilimod-induced vacuoles are associated with the endolysosomal system and support the possibility that impaired PIKfyve-dependent lysosomal trafficking contributes to hepatocyte vacuolation during ethanol-induced calcium dysregulation.

### TRPML1 activity regulates hepatocyte calcium dynamics, macrophage recruitment, and ethanol-associated vacuolation

Because PIKfyve generates PI(3,5)P₂, which directly activates the lysosomal calcium channel TRPML1 (Transient Receptor Potential Mucolipin 1) ^30,31^, we next investigated whether TRPML1-mediated calcium release contributes to the hepatocyte response to ethanol. To determine whether TRPML1 activation is sufficient to alter hepatocyte calcium dynamics, we treated 6 dpf *Tg(fabp10a:GCaMP6s)* larvae with 5 μM ML-SA1, a selective TRPML1 agonist ^32^. Vehicle-treated larvae displayed relatively stable GCaMP6s fluorescence, consistent with low basal calcium activity (**Supplementary Video 3**). In contrast, ML-SA1 treatment induced dynamic fluctuations in hepatocyte GCaMP6s fluorescence (**Figure 5A, Supplementary Video 4**). Quantification confirmed a significant increase in calcium oscillatory activity following ML-SA1 treatment compared with vehicle controls (p-value = 0.008, Mann–Whitney U test following Shapiro–Wilk normality assessment) (**Figure 5B**).

**Figure 5:**
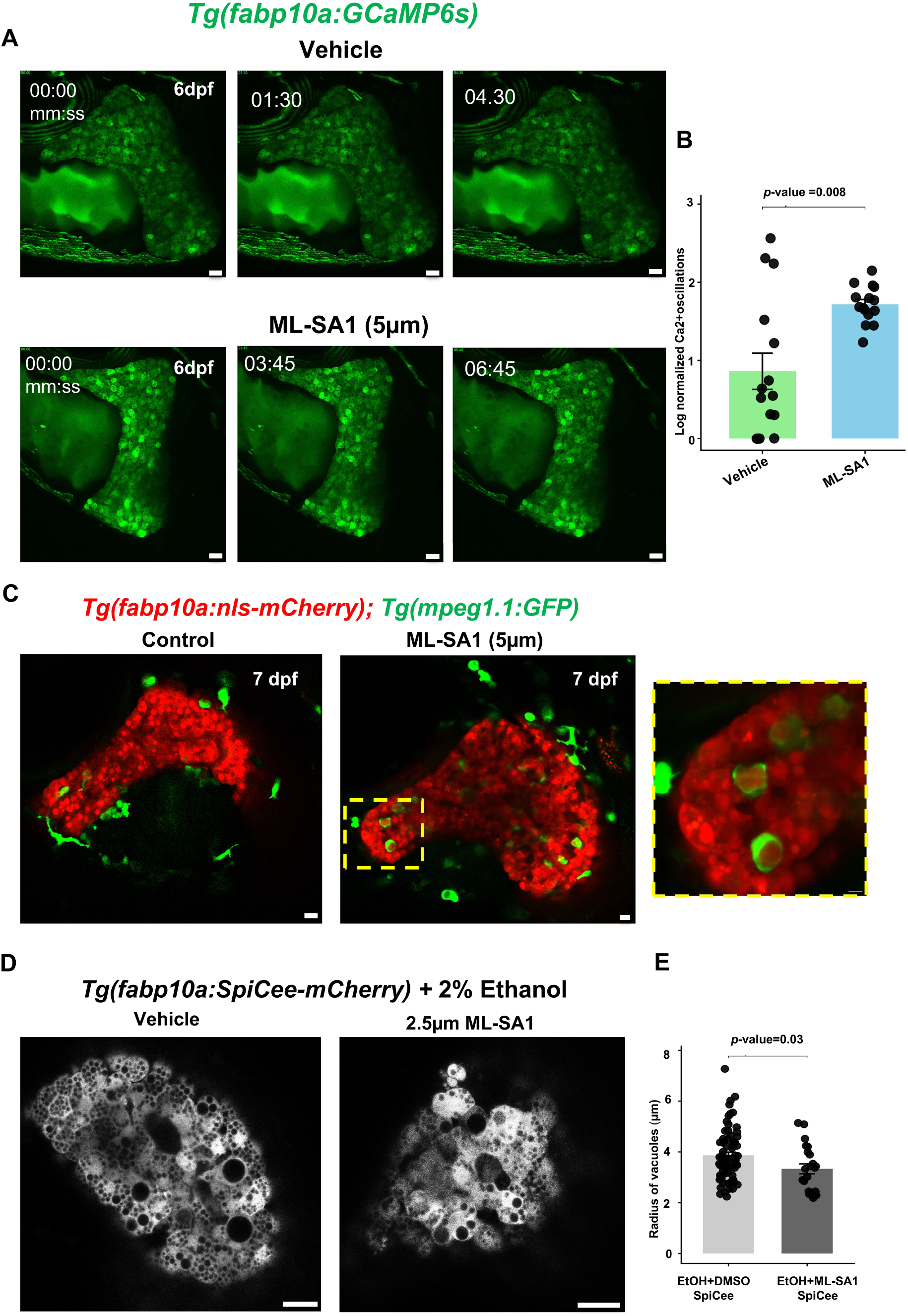
Activation of TRPML1 by ML-SA1 enhances calcium oscillatory activity in zebrafish hepatocytes. **(A)** Representative time-lapse confocal images of hepatocytes from 6 dpf *Tg(fabp10a:EGFP)* zebrafish larvae treated with vehicle as control and with 5 µM ML-SA1 (TRPML1 agonist). Time stamps indicate minutes:seconds (mm:ss). Scale bars 20 µm for all panels. **(B)** Quantification of normalized hepatocyte Ca²⁺ oscillations in vehicle and ML-SA1 treated larvae. ML-SA1 treatment significantly enhanced Ca²⁺ oscillatory activity compared with vehicle.Data are shown as mean ± SEM with individual biological replicates. Statistical significance was assessed using the Mann–Whitney U test after testing for normality using the Shapiro–Wilk test, p-value = 0.008. **(C)** ML-SA1 treatment (TRPML1 Agonist) induces macrophage accumulation within the liver. Representative confocal images of 7 dpf *Tg(fabp10a:nls-mCherry); Tg(mpeg1.1:GFP)* larvae showing hepatocyte nuclei (red) and macrophages (green). Compared with controls, ML-SA1-treated larvae displayed increased infiltration and accumulation of macrophages within the hepatic parenchyma. The boxed region is shown at higher magnification on the right, highlighting GFP-positive macrophages localized within hepatocytes. Scale bar 10um for Control and 5 µm for ML-SA1. **(D)** Confocal images of livers from *Tg(fabp10a:SpiCee-mCherry)* larvae exposed to 2% ethanol and treated with vehicle or ML-SA1 (2.5 μM). Scale bar = 20µm. Treatment with ML-SA1 slightly reduced vacuole size and severity, indicating partial rescue of the SpiCee- and ethanol-induced hepatic pathology. **(E)** Quantification of vacuolar size in SpiCee-expressing ethanol-treated livers. ML-SA1 treatment significantly reduced the radius of vacuolar structures compared with vehicle-treated controls. Each point represents an individual vacuole. Statistical significance was determined using a two-tailed Mann–Whitney U test following assessment of normality by the Shapiro–Wilk test. *p* -value= 0.03.

We next examined whether TRPML1 activation was sufficient to promote hepatic immune-cell recruitment. Double-transgenic *Tg(fabp10a:nls-mCherry)*;*Tg(mpeg1.1:EGFP)* larvae were treated with ML-SA1 and analysed at 7 dpf. In vehicle-treated animals, macrophages were located predominantly at the liver periphery. Following ML-SA1 treatment, increased numbers of GFP-positive macrophages were observed within the hepatic parenchyma (**Figure 5C**). These observations indicate that pharmacological activation of TRPML1 is sufficient to induce hepatocyte calcium activity and is associated with increased macrophage recruitment to the liver, mimicking the response of the zebrafish liver to alcohol exposure.

To test whether restoring lysosomal calcium release could reduce ethanol-associated vacuolation, *Tg(fabp10a:SpiCee-mCherry)* larvae were exposed to 2% ethanol and treated with either vehicle or 2.5 μM ML-SA1. Vehicle-treated larvae developed the prominent vacuolar phenotype observed previously in ethanol-exposed SpiCee-expressing hepatocytes (**Figure 5D**). ML-SA1 treatment produced a modest but significant reduction in vacuole radius (p-value = 0.03, Mann–Whitney U test) (**Figure 5D**), suggesting that enhancement of TRPML1-dependent calcium release can partially alleviate vacuolation caused by the combined effects of ethanol exposure and intracellular calcium buffering.

We then examined the complementary effect of TRPML1 inhibition using (1R,2R)-ML-SI3, a selective TRPML1 antagonist ^33,34^. Treatment of 6 dpf *Tg(fabp10a:nls-mCherry)* larvae with 5 μM (1R,2R)-ML-SI3 alone induced hepatocyte vacuolation in 4 of 10 animals, compared with 1 of 10 vehicle-treated controls (**Figure 6A**). Exposure to 1.5 % ethanol alone caused only limited vacuolation (1 of 10 larvae), whereas combined treatment with ethanol and (1R,2R)-ML-SI3 produced extensive vacuolation in all animals examined (10 of 10). The combined treatment also resulted in more numerous and larger vacuoles than either treatment alone (p-value = 0.0001, Fisher’s Exact Test), indicating that TRPML1 inhibition strongly enhances the hepatocyte response to ethanol. Here, we wish to note that increasing the ethanol concentration to 2 % in combination with (1R,2R)-ML-SI3 caused larval lethality, precluding morphological analysis under these conditions (data not shown). Further, a cis/trans mixture of ML-SI3 did not produce a detectable hepatic phenotype (data not shown), potentially because the inhibitory activity is primarily associated with the (−)-trans-stereoisomer (1R,2R) rather than the cis isomer mixture ^33^.

**Figure 6:**
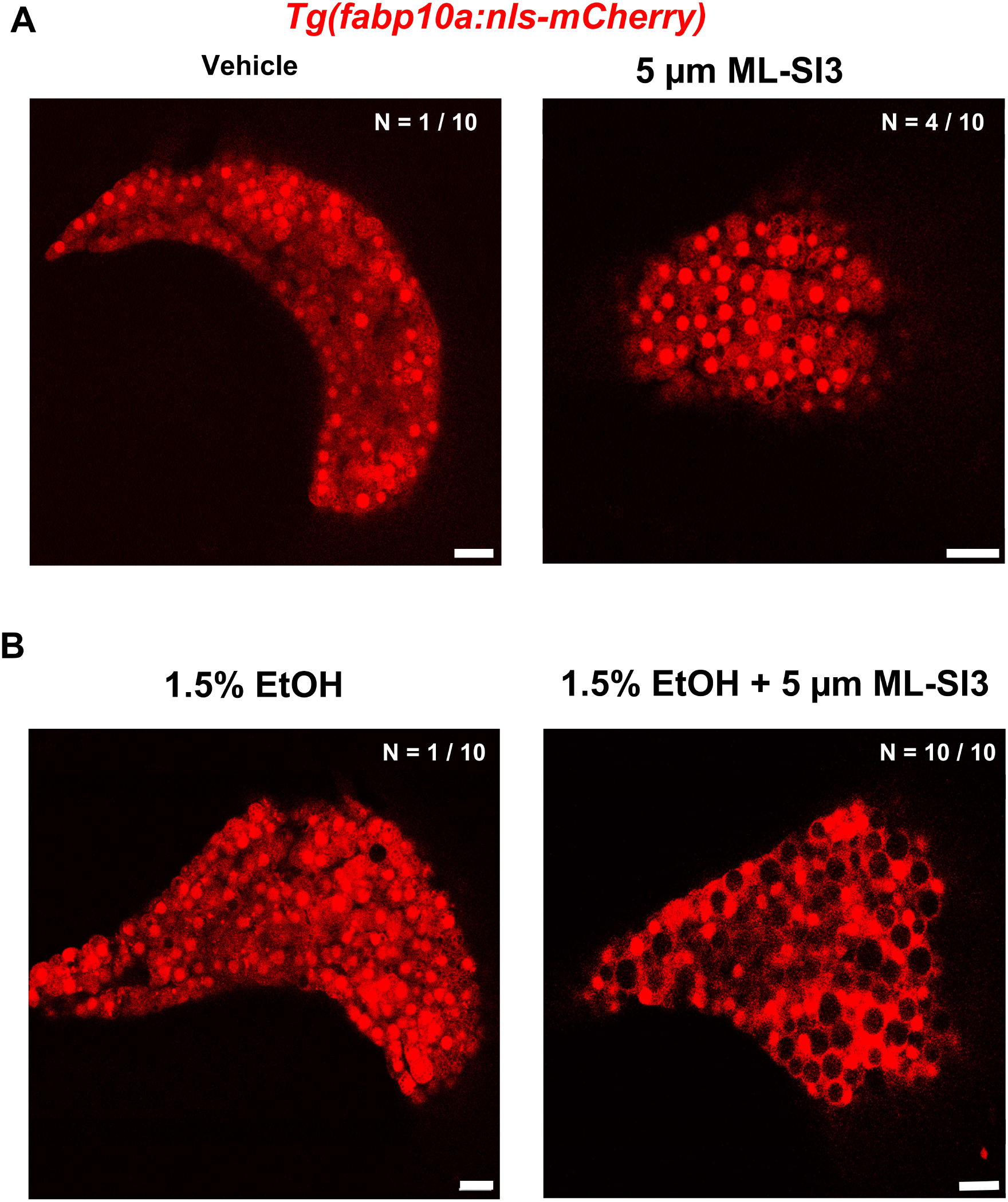
Trans ML-SI3 (TRPML1 antagonist) exposure induces a vacuolar phenotype in the zebrafish liver and exacerbates ethanol-induced hepatic vacuolation. **(A)** Representative confocal images of livers from 6 dpf *Tg(fabp10a:nls-mCherry)* larvae, in which hepatocyte nuclei are labeled with mCherry. Vehicle-treated larvae displayed normal liver morphology, whereas treatment with 5 μM ML-SI3 resulted in the formation of prominent intracellular vacuolar structures throughout the hepatic tissue. The incidence of the vacuolar phenotype is indicated in the image. Larvae were scored as vacuole-positive when more than 10 vacuoles per liver were detected. **(B)** Representative confocal images of livers from Tg(fabp10a:nls-mCherry) larvae exposed to 1.5% ethanol alone or in combination with 5 μM ML-SI3. Ethanol treatment induced hepatic vacuolation, while co-treatment with ML-SI3 further increased the severity of the vacuolar phenotype, characterized by larger and more abundant vacuolar structures. Phenotypic classification was performed using a threshold of >10 vacuoles per liver. Scale bars, 20 μm for all panels.

Together, these complementary pharmacological experiments support a relationship between TRPML1 activity and hepatocyte vacuolation. TRPML1 activation induces calcium oscillations, promotes macrophage recruitment, and partially reduces ethanol-associated vacuolation, whereas TRPML1 inhibition induces vacuolation and markedly exacerbates the effect of ethanol. These findings implicate the PIKfyve–PI(3,5)P₂–TRPML1 pathway as an important regulator of lysosomal calcium signalling and hepatocyte homeostasis during ethanol exposure.

### Ethanol exposure and calcium buffering promote intracellular aggregate accumulation in pancreatic acinar cells

Ethanol induced asynchronous calcium oscillations in pancreatic acinar cells (**Figure 3**) and altered the expression of calcium-signalling genes in this population (**Figure 2D**). We therefore asked whether these changes were associated with downstream cellular abnormalities *in vivo*. To address this, we examined 6 dpf *Tg(ela3l:dTomato)* zebrafish larvae, in which dTomato is specifically expressed in pancreatic acinar cells. Under control conditions, dTomato fluorescence was distributed relatively uniformly throughout the acinar tissue, with only occasional punctate structures (**Figure 7A**). Following exposure to 2% ethanol for 19–22 hours, acinar cells developed multiple bright intracellular puncta (**Figure 7A**), resulting in a significant increase in aggregate-positive area (p-value = 0.0184) (**Figure 7C**).

**Figure 7:**
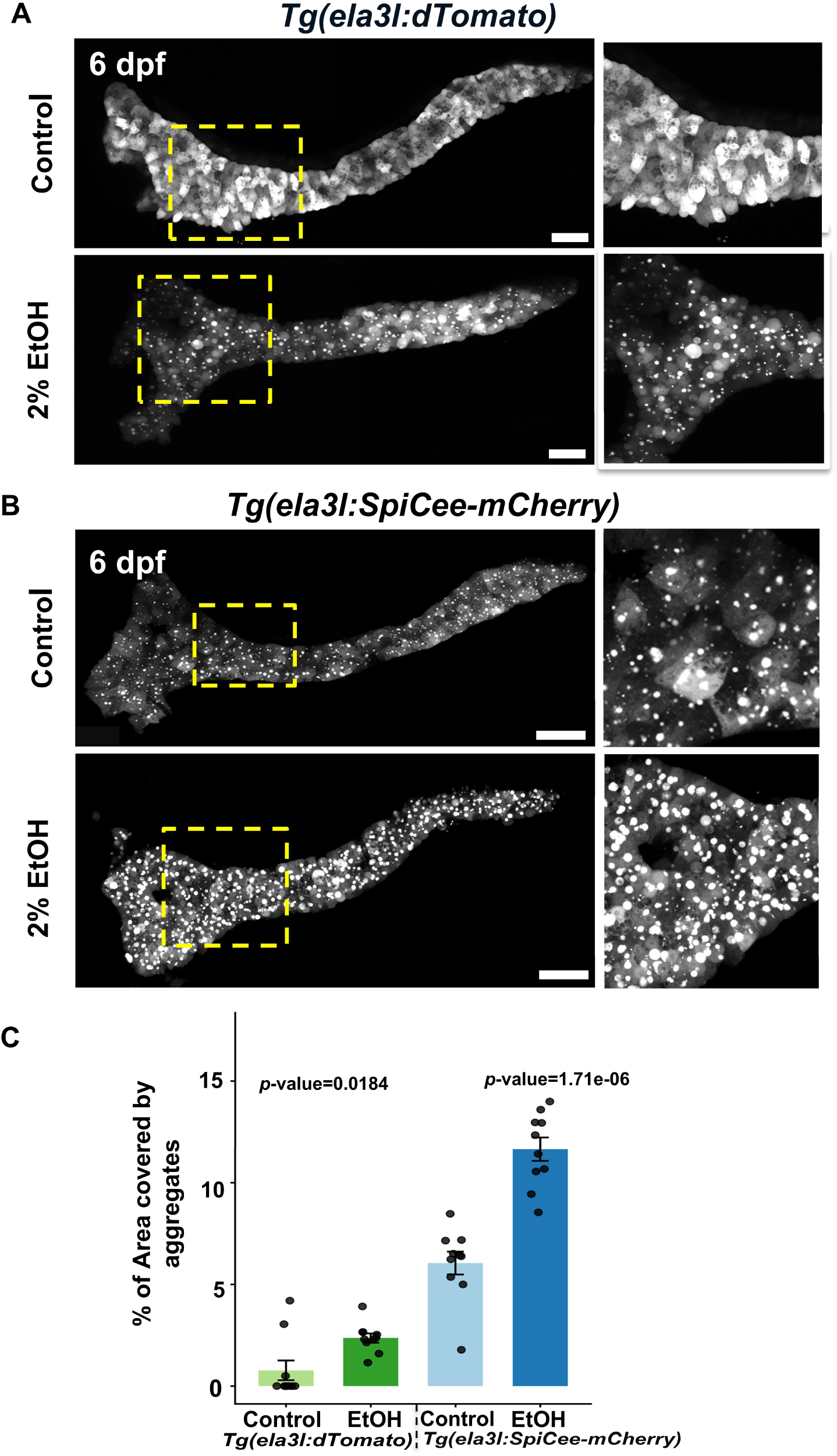
Ethanol exposure promotes intracellular aggregate accumulation in pancreatic acinar cells of zebrafish larvae. **(A)** Confocal images of pancreatic acinar cells from 6 dpf *Tg(ela3l:dTomato)* zebrafish larvae under control and 2 % ethanol (EtOH) treatment conditions for 19 hrs-22 hrs.Yellow dashed boxes indicate regions shown at higher magnification in the right panels. Scale bar 50 μm. **(B)** Confocal images of pancreatic acinar cells from 6 dpf *Tg(ela3l:SpiCee-mCherry)* zebrafish larvae expressing the genetically encoded SpiCee which causes calcium buffering under control and 2 % ethanol (EtOH) treatment conditions. Yellow dashed boxes indicate regions shown at higher magnification in the right panels. Scale bars 50 μm. **(C)** Quantification of aggregate-positive area in pancreatic acinar cells of *Tg(ela3l:dTomato)* and Tg(ela3l:SpiCee-mCherry) larvae following ethanol exposure. Ethanol treatment significantly increased aggregate accumulation in both reporter lines. Data are shown as mean ± SEM; each dot represents one biological replicate. Statistical significance was assessed by pairwise unpaired comparisons following Shapiro–Wilk normality testing. The Tg(ela3l:dTomato) Control versus EtOH comparison was analyzed using a two-tailed Wilcoxon rank-sum test (*p* = 0.0184), whereas the Tg(ela3l:SpiCee-mCherry) Control versus EtOH comparison was analyzed using a two-tailed Welch’s *t*-test (*p* = 1.71 × 10⁻⁶).

To determine whether calcium signalling was required to maintain acinar-cell homeostasis, we next examined *Tg(ela3l:mCherry-SpiCee)* larvae expressing the genetically encoded calcium buffer SpiCee specifically in acinar cells. Even in the absence of ethanol, SpiCee-expressing larvae displayed a higher basal abundance of fluorescent aggregates than controls (**Figure 7A-C**), indicating that sustained calcium buffering alone disrupts intracellular protein or vesicle handling. Exposure of SpiCee-expressing larvae to 2% ethanol further increased aggregate accumulation beyond that observed with either ethanol exposure or calcium buffering alone (p-value = 1.71 × 10⁻⁶, two-tailed Welch’s *t*-test) (**Figure 7C**). Importantly, unlike the vacuolar phenotype observed in hepatocytes (**Figure 4A**), calcium disruption in acinar cells was associated predominantly with intracellular aggregate accumulation, suggesting distinct secretory and proteostatic demands of this cell type.

### PICK1 inhibition exacerbates ethanol-induced aggregate accumulation in pancreatic acinar cells

Because PIKfyve was implicated in the hepatocyte response, we first asked whether inhibition of this pathway produced a comparable phenotype in pancreatic acinar cells. Treatment of *Tg(ela3l:dTomato)* larvae with 0.5 μM apilimod induced extensive intracellular vacuolation, whereas vehicle-treated larvae retained normal acinar morphology (**Supplementary Figure 4**). Thus, disruption of PIKfyve-dependent trafficking affects both hepatocytes and acinar cells, although the dominant ethanol-associated phenotype in acinar cells was aggregate accumulation rather than vacuolation.

Single-cell transcriptomic analysis identified significant upregulation of *pick1* in ethanol-exposed pancreatic acinar cells (**Figure 2D**), prompting us to investigate its functional relevance. *pick1* encodes Protein Interacting with C Kinase 1 (PICK1) ^35^, PDZ domain-containing scaffold protein involved in membrane organization and vesicular trafficking. PICK1 also has a documented relationship with calcium signalling: calcium can bind to PICK1 and regulate its membrane-associated trafficking functions, leading to its description as a calcium sensor in activity-dependent cargo trafficking ^36,37^. Its induction following ethanol exposure could therefore reflect a compensatory response that helps preserve intracellular trafficking.

To test this, we pharmacologically inhibited PICK1 using FSC231, a well established inhibitor of PICK1 PDZ domain ^38^. In 6 dpf *Tg(ela3l:dTomato)* zebrafish larvae, under basal conditions, 10µM FSC231 treatment did not significantly alter acinar-cell morphology or aggregate-positive area compared with vehicle-treated controls (**Figure 8A, B**). Thus, PICK1 inhibition alone was insufficient to induce a detectable phenotype under the conditions examined.

**Figure 8:**
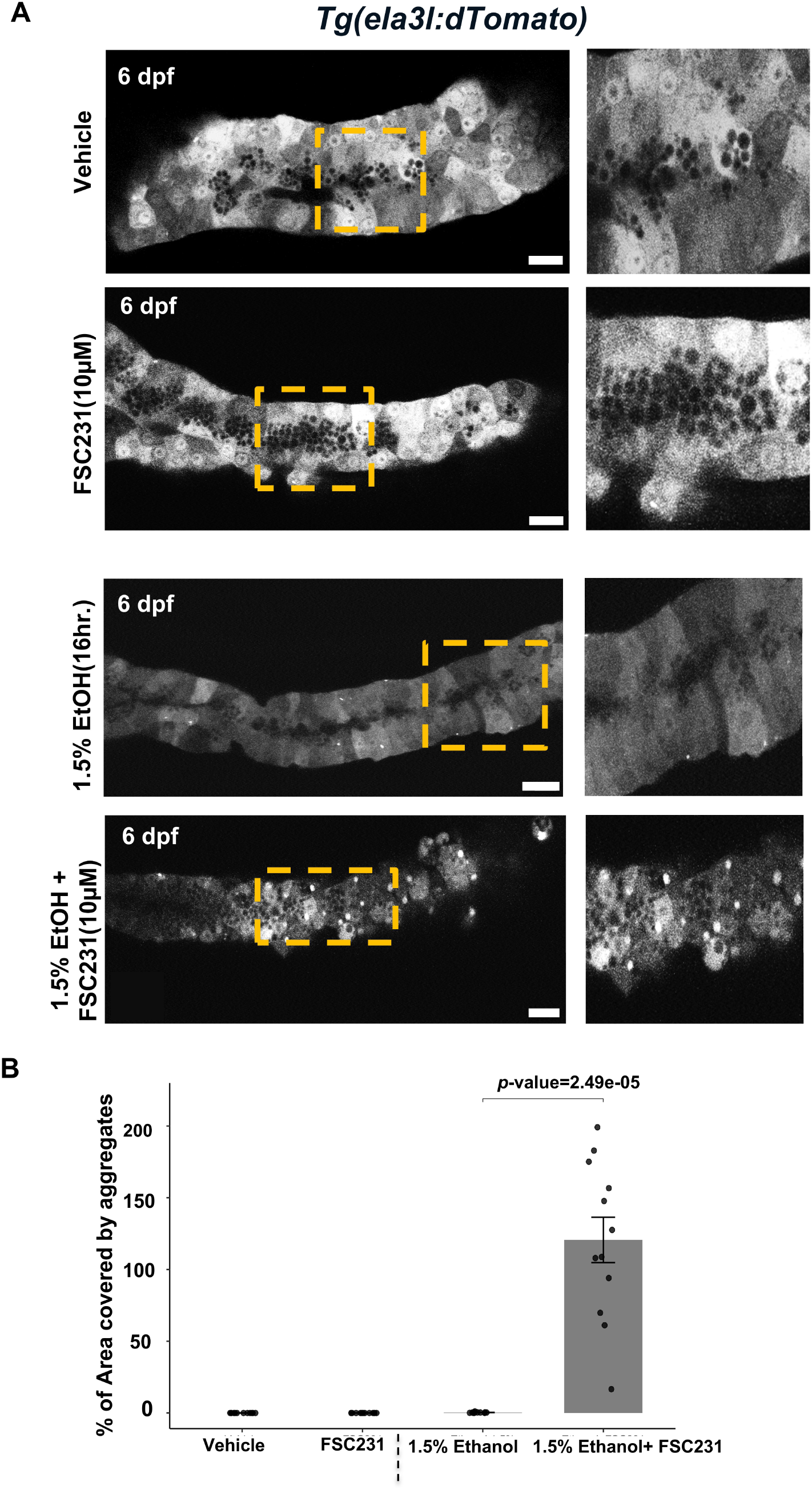
PICK1 inhibition exacerbates aggregate accumulation in ethanol-treated pancreatic acinar cells. **(A)** Confocal images of pancreatic acinar cells from 6 dpf *Tg(ela3l:dTomato)* zebrafish larvae treated with vehicle (control) or the PICK1 inhibitor FSC231 under basal conditions; or exposed to 1.5% ethanol (EtOH) in the absence or presence of FSC231. Yellow dashed boxes indicate regions shown at higher magnification in the right panels. Scale bars: 50μm. **(B)** Quantification of the percentage area occupied by aggregates in pancreatic tissue from *Tg(ela3l:dTomato)* zebrafish larvae under vehicle, FSC231, ethanol (1.5%), and ethanol + FSC231 treatment conditions. Data are presented as mean ± SEM, with each dot representing one biological replicate. Statistical analysis was performed using pairwise comparisons following Shapiro–Wilk normality testing. Vehicle vs FSC231: not significant; Ethanol vs Ethanol + FSC231: *p*-value = 2.49 × 10⁻⁵(two tailed Wilcoxon rank-sum test).

We next exposed larvae to 1.5% ethanol either alone or in combination with FSC231(10uM). Ethanol exposure alone produced only limited aggregate accumulation. In contrast, combined treatment with ethanol and FSC231 caused a marked increase in intracellular aggregates throughout the acinar tissue (**Figure 8A, B**). Quantification confirmed a significant increase in aggregate-positive area following combined treatment compared with ethanol alone (p-value = 2.49 × 10⁻⁵, two tailed Wilcoxon rank-sum test)

These findings suggest that PICK1 is not required for basal acinar-cell morphology but becomes important during ethanol-induced stress. The strong enhancement of aggregate accumulation following PICK1 inhibition is consistent with a protective or compensatory role for PICK1-dependent trafficking in maintaining acinar-cell proteostasis during alcohol exposure.

## Discussion

Our findings support a model in which calcium signalling is an active determinant of cellular adaptation to alcohol stress rather than merely a downstream consequence of injury. In both hepatocytes and pancreatic acinar cells, broad suppression of calcium dynamics worsened cellular pathology, indicating that calcium flux is required to maintain homeostasis during ethanol exposure. However, the consequences of calcium disruption differed between the two tissues, suggesting that a shared upstream stress response is interpreted through cell-type-specific organelle and trafficking networks.

In hepatocytes, the predominant consequence of calcium buffering was the accumulation of large intracellular vacuoles (**Figure 4A**), placing the endolysosomal system at the centre of the adaptive response. This phenotype was reproduced by inhibition of PIKfyve (**Figure 4B**), a lipid kinase that generates PI(3,5)P₂ and thereby regulates endolysosomal membrane dynamics and trafficking. *In vivo* evidence from zebrafish strongly supports this interpretation. *pikfyve* mutants develop severe larval-stage defects and die by 9 dpf, but before lethality they accumulate prominent vacuoles in the lens that produce an early-onset cataract phenotype resembling congenital cataract in humans ^39^. These vacuoles were identified as Rab7- and LC3-positive amphisomes, consistent with defective late endosomal and autophagic processing. Similarly, loss-of-function mutations in *fig4a*, which encodes a component of the PIKfyve complex required for PI(3,5)P₂ homeostasis, cause robust hepatic vacuolation associated with abnormal lysosomal storage and accumulation of autophagic intermediates ^26^. Together, these studies provide direct in vivo evidence that disruption of the PIKfyve–PI(3,5)P₂ pathway is sufficient to impair lysosomal processing in zebrafish tissues, including hepatocytes.

Notably, *pikfyve* was not transcriptionally upregulated in either the single-cell atlas generated here or in bulk RNA-seq from our previous study ^11^, despite the strong phenotype caused by inhibition of PIKfyve. This suggests that pathway engagement may occur through post-transcriptional regulation or increased functional demand rather than changes in gene expression, and highlights tissue-specific calcium buffering as a complementary strategy for revealing stress-sensitive cellular dependencies that may not be evident from transcriptomic analysis alone.

TRPML1 provides a mechanistic link between PIKfyve activity and lysosomal calcium release. The similarity between calcium-buffering-induced vacuolation and TRPML1 inhibition (**Figure 6**), together with the partial reduction of vacuolation following TRPML1 activation (**Figure 5D**), suggests that lysosomal calcium release supports the maturation and resolution of vesicular compartments during ethanol stress. This interpretation is consistent with published evidence that TRPML1 regulates lysosome size through calcium-dependent calmodulin activation and promotes autophagosome–lysosome fusion, lysosomal acidification, and recruitment of fusion-associated SNARE proteins ^40,41^. It is also supported by the enlarged lysosomes and accumulation of autophagic material reported in skeletal muscle of *mcoln1a/mcoln1b* double-mutant zebrafish ^42^. Our findings extend these observations to hepatocytes and suggest that ethanol exposure increases cellular dependence on TRPML1-mediated lysosomal processing.

The strong interaction between ethanol and TRPML1 inhibition suggests that TRPML1 becomes particularly important when lysosomal demand is increased. Hepatocytes may compensate for partial TRPML1 inhibition under basal conditions, whereas ethanol-induced trafficking, organelle damage, and degradative burden expose a latent dependence on lysosomal calcium release.

TRPML1 activation nevertheless also increased hepatic macrophage recruitment (**Figure 5C**), indicating that lysosomal calcium signalling may have both adaptive and pro-inflammatory consequences. Calcium release within a physiological range may support membrane trafficking and degradative function, whereas excessive or prolonged activation may promote stress signalling, damage-associated cues, or sterile inflammation. Such a threshold-dependent response provides a possible explanation for how transient alcohol-induced stress can be resolved, while repeated or sustained exposure progressively shifts the tissue towards inflammation and injury.

In pancreatic acinar cells, calcium buffering instead promoted intracellular aggregate accumulation (**Figure 7C**). This difference likely reflects the exceptional secretory burden of acinar cells, which depend on tightly regulated calcium signals for protein folding, zymogen trafficking, and secretion. Previous studies showed that non-oxidative ethanol metabolites induce IP₃ receptor-dependent calcium release, mitochondrial dysfunction, ATP depletion, and sustained calcium toxicity in acinar cells ^43,44^. Our findings complement this work by showing that broad calcium suppression is also detrimental. Residual or oscillatory calcium activity may therefore be required to sustain proteostasis during ethanol stress. The enhancement of aggregation following PICK1 inhibition (**Figure 8**) further suggests that PICK1 contributes to a compensatory trafficking programme that limits the accumulation of improperly processed proteins.

The distinct cellular consequences observed in hepatocytes and pancreatic acinar cells suggest that a single therapeutic strategy may not be sufficient to protect both organs in alcohol-related disease. Although calcium dysregulation may represent a shared upstream feature, the dominant downstream vulnerabilities appear to differ between liver and pancreas. These findings raise the possibility that tissue-tailored interventions, or carefully designed multi-pathway approaches, may ultimately be required when both organs are affected. However, several limitations should be considered. The acute larval zebrafish model does not reproduce the duration or systemic complexity of chronic human alcohol-associated disease. SpiCee buffers calcium broadly and cannot resolve the contributions of individual intracellular stores, while pharmacological manipulation of PIKfyve, TRPML1, and PICK1 may include off-target effects. Nevertheless, the study supports a framework in which alcohol engages a common calcium-dependent stress response, while tissue-specific trafficking and organelle demands determine whether its disruption manifests as lysosomal dysfunction, proteotoxic stress, or inflammation.

## Methods

### Zebrafish lines and husbandry

Wild-type and transgenic zebrafish from the outbred AB strain were utilized in all experiments. All zebrafish husbandry and experimental procedures for transgenic lines were conducted in accordance with institutional and national ethical and animal welfare guidelines and regulations, which were approved by the Ethical Committee for Animal Welfare (CEBEA) from the Université Libre de Bruxelles (protocols 864N, 865N, 877N, 881N, 882N) and Shiv Nadar Institution of Eminence (SNIoE/IAEC/2026/1/004).

In this study, the following published transgenic lines were used: *Tg(fabp10a:FLAG-SpiCee-mCherry; cryaa:mCherry)^ulb^*^16^, Tg(fabp10a:GCaMP6s; cryaa:mCherry)^ulb^^15^ ^11,23^, *Tg(mpeg1.1:EGFP)*^gl^^22^ ^45^, *Tg(fabp10a:nls-mCherry)*^mss^^4^ ^46^, and *Tg(fabp10a:EGFP)*^as^^3^ ^47^.

The following lines were newly generated by the I-SceI system: *Tg(ela3l:lox2272-loxp-dTomato-stop-lox2272-mCerulean-stop-loxp-EYFP; cryaa:mCherry)^ulb^*^53^ abbreviated as *Tg(ela3l:dTomato)*, Tg(ela3l:GCaMP6s; cryaa:mCherry)^ulb^^54^ and *Tg(fabp10a:FLAG-SpiCee-mCherry; cryaa:mCherry)^ulb^*^55^.

### Generation of the Tg(ela3l:dTomato)^ulb^^53^, Tg(ela3l:GCaMP6s)^ulb^^54^ and Tg(ela3l:SpiCee-mCherry)^ulb^^55^ lines

The 2420-bp sequence upstream of the *ela3l* gene was used to generate transgenic lines targeting pancreatic acinar cells ^48^. The sequence was retrieved from Ensembl using the GRCz11 zebrafish genome assembly and synthesized by GenScript Biotech in the pUC57 vector. The synthesized fragment was flanked by SacI at the 5′ end and AscI and NheI sites at the 3′ end. To generate the *ela3l* construct, the promoter was excised using SacI and AscI and cloned into the ins:lox2272-loxp-dTomato-stop-lox2272-mCerulean-stop-loxp-EYFP; cryaa:mCherry (ins:Brainbow1.0L; cryaa:mCherry) plasmid ^49^, replacing the *ins* promoter. The resulting construct ela3l:lox2272-loxp-dTomato-stop-lox2272-mCerulean-stop-loxp-EYFP; cryaa:mCherry (ela3l:BB1.0L; cryaa:mCherry) was designated ela3l:dTomato because dTomato is the default fluorescent protein expressed by the Brainbow1.0L cassette ^50^. Similarly, the *ela3l* promoter was cloned using SacI and NheI into the ins:GCaMP6s; cryaa:mCherry plasmid ^49^ using SacI and SpeI and into fabp10a:FLAG-SpiCee-mCherry; cryaa:mCherry ^11^ plasmid using SacI and NheI.

To generate transgenic lines, a solution containing the 20 ng/µl of the construct was mixed with I-SceI meganuclease enzyme and injected into one-cell stage embryos to facilitate transgenesis. Transgenic founders were identified and maintained using the eye marker expression.

### Single-cell RNA sequencing and data analysis

Single-cell suspensions and transcriptomic libraries were prepared exactly as previously described ^51^. Briefly, the dissected digestive organs were dissociated in TrypLE at 37°C, filtered through a 40-µm cell strainer, and stained with Calcein violet to identify viable cells. Live Calcein-positive cells were sorted using a BD FACSAria II with a 100-µm nozzle, and approximately 50,000 cells were collected per condition. The collection tubes were coated with 1% BSA in PBS to lower cells from sticking to the tubes ^52^. Single-cell RNA-sequencing libraries were generated using the 10x Genomics Chromium Single Cell 3′ v3 platform, targeting approximately 10,000 cells per sample, and sequenced on an Illumina NextSeq 550. Reads were aligned to the Ensembl GRCz11 zebrafish genome, and gene-expression matrices were generated using Cell Ranger v7.1.0.

Downstream analysis was performed in Seurat ^53^ following the standard workflow. Cells with fewer than 500 or more than 8,000 detected genes, or with more than 15% mitochondrial transcripts, were excluded. Data were log-normalized, scaled, and analyzed by principal component analysis and UMAP. Clustering was performed at a resolution of 0.8, and cell types were assigned based on established marker genes. Differential gene-expression analysis between control and ethanol-treated cells was performed using the Wilcoxon rank-sum test.

### Pharmacological Treatments

Apilimod (Sigma-Aldrich, SML2974), ML-SA1 (Tocris, SML0627), (1R,2R)-ML-SI3 (MedChemExpress, HY-134819A), and FSC231 (MedChemExpress, HY-117772) were prepared as 1000× stock solutions in DMSO and stored at −20 °C or −80 °C. Working solutions were obtained by dilution in embryo medium and added directly to the embryos. Apilimod incubations were performed in the dark. Drug concentrations and treatment durations are specified in the corresponding Results sections and figure legends. For live-imaging experiments, the respective compounds were maintained in the medium throughout image acquisition. Equivalent concentrations of DMSO were used as vehicle controls.

### Imaging and image analysis

For *in vivo* confocal imaging, larvae were anesthetized with 0.02% tricaine (MS-222) (Sigma-Aldrich, E10521) or immobilized with 4 mM (+)-tubocurarine (for GCaMP6s imaging) (Sigma-Aldrich, 93750), immobilized in 1% Low-Melt Agarose (Lonza, 50080), and imaged in glass-bottomed dish FluoroDish™ (WPI, FD3510-100) using a Zeiss LSM 780 confocal microscope. The imaging frame was set to 1024 × 1024 pixels. Samples were excited at 488 nm for GCaMP6s / EGFP, and 543 nm for mCherry / dTomato with fluorescence collected in the respective ranges of 497-532 nm and 550-633 nm.

For calcium imaging, Z-stack time-lapse images were acquired at 45-s intervals. Maximum-intensity projections were generated in Fiji, and individual cells were manually defined as regions of interest. Mean GCaMP6s fluorescence intensity was extracted for each cell and analyzed in R ^54^. Fluorescence values were normalized to the local minimum, and calcium oscillations were identified as local intensity maxima. Oscillation numbers were summed per image and, where indicated, normalized to tissue volume and imaging duration.

For Nile Red staining, a 500 µg/ml stock solution was prepared in acetone. Larvae were incubated for 1 h at 28°C in a freshly prepared 1:300 dilution of Nile Red in E2 medium, protected from light, and washed before imaging. EGFP-positive hepatocytes and Nile Red-labelled neutral lipids were imaged spectrally, as published previously ^11,23^. Lipid droplets within the hepatocyte region were manually identified and quantified in Fiji, and the corresponding liver area was measured for normalization.

Animals were incubated in 1:100 Lysotracker Green DND-26 (ThermoFisher, L7526) or Lysotracker Red DND-99 (ThermoFisher, L7528) dissolved in embryo (E2) medium in the dark at 28°C. Five larvae were treated in 1 mL of medium. After 1.5 hours of incubation, imaging was initiated. To avoid cytotoxicity, imaging was performed during the next 30 min after the incubation time in lysotracker. As control, DMSO-treated animals were used.

Hepatocyte vacuoles were identified in confocal images as non-fluorescent, optically empty spaces greater than 4 µm in diameter within the EGFP-positive liver parenchyma. Animals were scored according to the presence or absence of one or more vacuoles.

Pancreatic acinar cell aggregation was assessed using maximum-intensity projections of confocal Z-stacks. Upper and lower intensity thresholds were set for each image, and the thresholded images were converted into binary masks. A watershed algorithm was applied to separate adjacent particles. The pancreatic region was manually defined as a region of interest (ROI), and its total area was measured. Particles within the ROI were quantified using the Analyze Particles function in Fiji, and the percentage of particle-positive pancreatic area was calculated as: Particle-positive area (%) = (total particle area / total pancreatic area) × 100.

Abnormally concentrated or clustered dTomato/mCherry-positive structures within the exocrine pancreas were classified as acinar aggregates. Animals were also scored according to the presence or absence of these aggregates..

### Statistical analysis

Statistical analysis was performed using R version 4.2.0 ^55^. Data distributions were assessed using a normality test, and appropriate parametric or non-parametric statistical tests were applied accordingly. Comparisons involving multiple groups were performed using two-way ANOVA followed by an appropriate multiple-comparisons test when the assumptions for parametric analysis were met. Fisher’s exact test was used to compare the number or proportion of animals displaying a phenotype between groups. The specific statistical test used for each analysis is indicated in the corresponding figure legend. No data were excluded from the analyses, and blinding was not performed during data analysis.

## Data Availability

The raw files and raw count table from deep sequencing can be accessed at Gene Expression Omnibus (GEO) with accession number GSE316507 with the reviewer token mhwtuygcrjghxsp.

Raw images and image analysis files are available upon request to the corresponding author. Source data are provided with this paper.

## Acknowledgements

We thank the members of IRIBHM and SNU Fish Facility, M Martens and JM Vanderwinden from the Light Microscopy Facility for technical assistance at ULB. The work was supported by FNRS grants 40006730 (ASP) to SEE, and 40005588 (MISU-PROL), 40013427 (CDR), 40027730 (CDR) and 40020360 (PDR) to SPS, and funding from Université libre de Bruxelles, Jaumotte-Demoulin Foundation, Ramalingaswami Re-entry Fellowship (BT/RLF/Re-entry/03/2023) from Department of Biotechnology (DBT), India, and Anusandhan National Research Foundation (ANRF) Advanced Research Grant (ANRF/ARG/2025/003178/LS) to SPS.

## Author Contributions

KG: investigation, visualization, methodology, writing—original draft, review, and editing. MPM: conceptualization, investigation, visualization, methodology

SEE, AT: investigation, methodology and resources.

RM: supervision, methodology, and writing—original draft, review and editing.

SPS: conceptualization, supervision, funding acquisition, project administration, and writing— original draft, review, and editing.

