## Supplemental Figures 1-4, Table 1, Video 1-4 for "Alcohol-Evoked Calcium signalling Drives Distinct Responses in Zebrafish Hepatocytes and Pancreatic Acinar Cells": Supplementary Figures.pdf

### Table of Contents

Supplementary Figure 1: Single-cell transcriptomic analysis of intestinal, renal, stromal, immune, and proliferating cell populations following ethanol exposure. 4

Supplementary Figure 2: Ethanol exposure induces stress-related transcriptional signature in hepatocytes. 6

Supplementary Figure 3: Ethanol exposure induces macrophage infiltration into the pancreas. 7

Supplementary Figure 4: Pharmacological inhibition of PIKfyve induces vacuolar phenotype in pancreatic acinar cells. 8

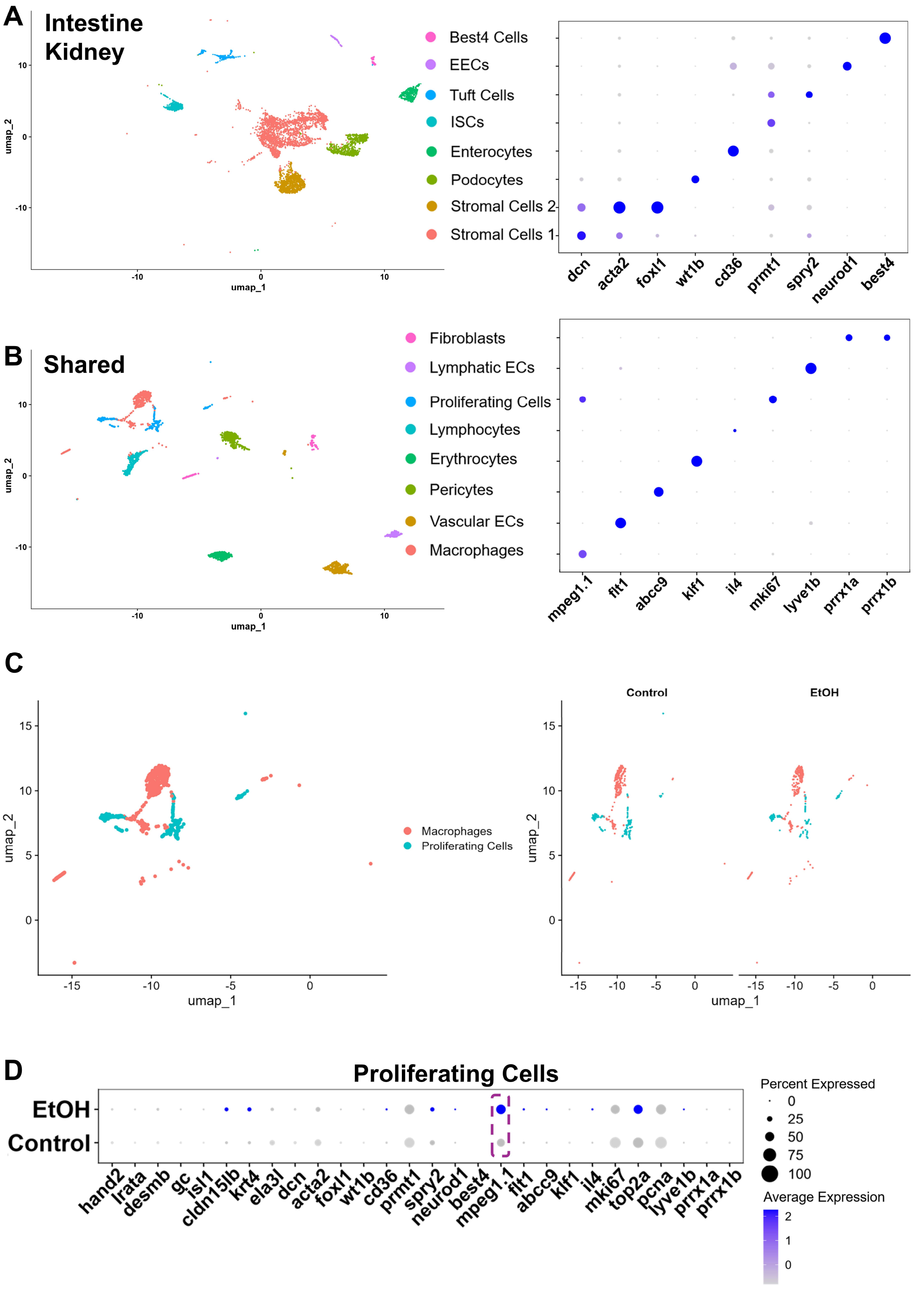
A
Intestine
Kidney
Best4 Cells
EECs
Tuft Cells
ISCs
Enterocytes
Podocytes
Stromal Cells 2
Stromal Cells 1
dcn
acta2
foxl1
wt1b
cd36
prmt1
spry2
neurod1
best4
umap_1
umap_2
B
Shared
Fibroblasts
Lymphatic ECs
Proliferating Cells
Lymphocytes
Erythrocytes
Pericytes
Vascular ECs
Macrophages
mpeg1.1
flt1
abcc9
klf1
il4
mki67
lyve1b
prrx1a
prrx1b
C
Macrophages
Proliferating Cells
Control
EtOH
D
Proliferating Cells
EtOH
Control
hand2
lrata
desmb
gc
isl1
cldn15lb
krt4
ela3l
dcn
acta2
foxl1
wt1b
cd36
prmt1
spry2
neurod1
best4
mpeg1.1
flt1
abcc9
klf1
il4
mki67
top2a
pcna
lyve1b
prrx1a
prrx1b
Percent Expressed
0
25
50
75
100
Average Expression
2
1
0

**Supplementary Figure 1: Single-cell transcriptomic analysis of intestinal, renal, stromal, immune, and proliferating cell populations following ethanol exposure.**

**(A)** Sub-clustering analysis of intestine- and kidney-derived cells. Corresponding dot plots show expression patterns of marker genes used for cell-type assignment.

**(B)** Characterization of shared cell populations present across multiple digestive organs. Dot plots depict representative marker genes used for annotation of each shared cell population.

**(C)** (Left) UMAP of the shared-cell compartment with macrophages and proliferating cells labelled. (Right) Split UMAP projections showcasing control and ethanol-treated (EtOH) samples.

**(D)** Dot plot showing the expression of lineage-associated and proliferation-related marker genes under control and ethanol-treated conditions. The macrophage marker, mpeg1.1, is enriched in the EtOH sample.

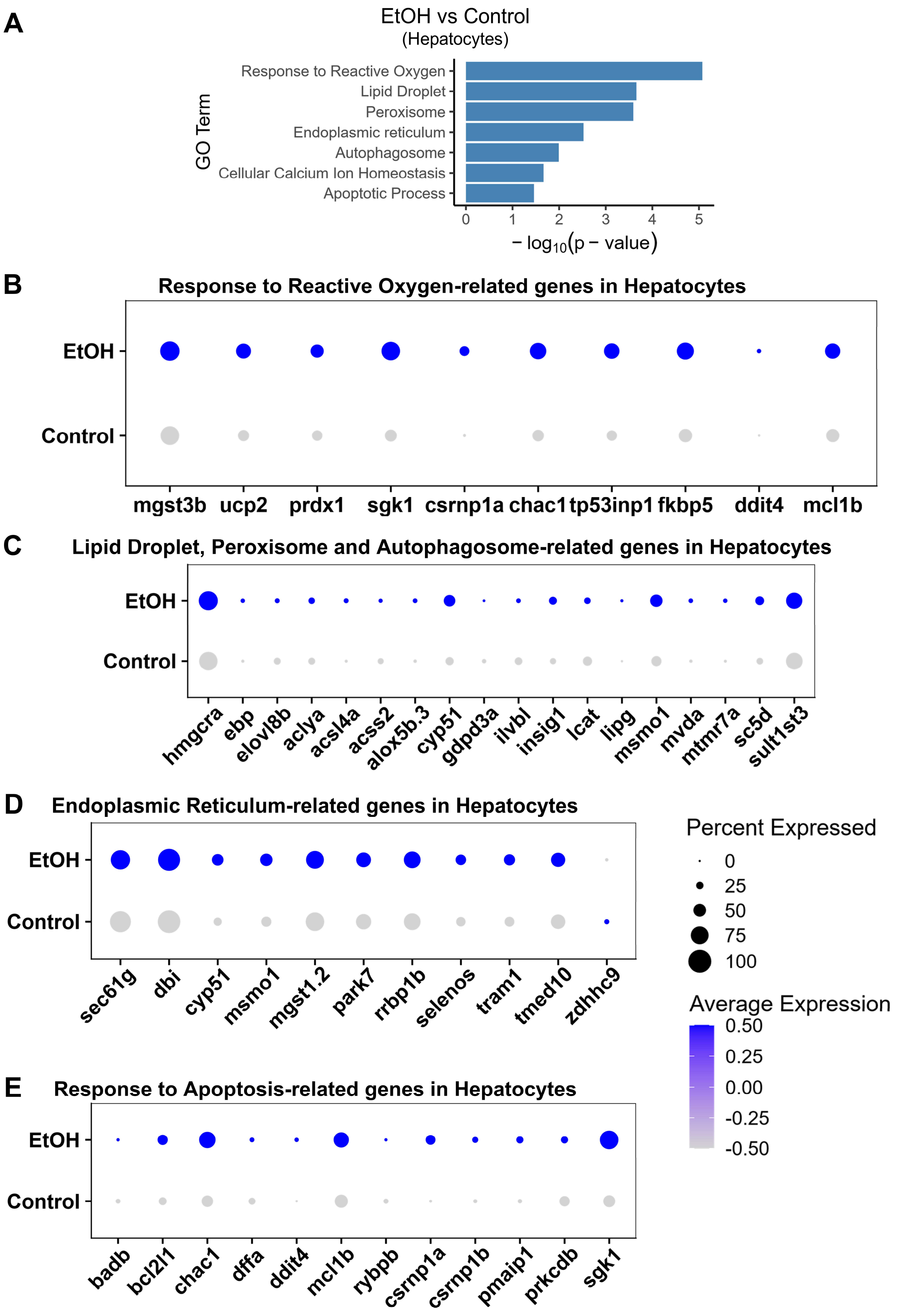
A
EtOH vs Control
(Hepatocytes)
GO Term
Response to Reactive Oxygen
Lipid Droplet
Peroxisome
Endoplasmic reticulum
Autophagosome
Cellular Calcium Ion Homeostasis
Apoptotic Process
0 1 2 3 4 5
− log10(p − value)
B
Response to Reactive Oxygen-related genes in Hepatocytes
EtOH
Control
mgst3b ucp2 prdx1 sgk1 csrnp1a chac1 tp53inp1 fkbp5 ddit4 mcl1b
C
Lipid Droplet, Peroxisome and Autophagosome-related genes in Hepatocytes
EtOH
Control
hmgcra ebp elovl8b aclya acsl4a acss2 alox5b.3 cyp51 gdpd3a ilvbl insig1 lcat lipg msmo1 mvda mtmr7a sc5d sult1st3
D
Endoplasmic Reticulum-related genes in Hepatocytes
EtOH
Control
sec61g dbi cyp51 msmo1 mgst1.2 park7 rrbp1b selenos tram1 tmed10 zdhhc9
Percent Expressed
0
25
50
75
100
Average Expression
0.50
0.25
0.00
-0.25
-0.50
E
Response to Apoptosis-related genes in Hepatocytes
EtOH
Control
badb bcl2l1 chac1 dffa ddit4 mcl1b rybpb csrnp1a csrnp1b pmaip1 prkcdb sgk1

**Supplementary Figure 2: Ethanol exposure induces stress-related transcriptional signature in hepatocytes.**

**(A)** Gene Ontology (GO) enrichment analysis of differentially expressed genes identified in hepatocytes following ethanol (EtOH) exposure compared with controls. Significantly enriched biological processes and cellular pathways included responses to reactive oxygen species (ROS), lipid droplet formation, peroxisome-associated pathways, endoplasmic reticulum (ER) function, autophagy-related processes, cellular calcium ion homeostasis, and apoptosis. Bar length represents enrichment significance expressed as $-\log_{10}$(p-value).

Dot plot showing expression of ROS-responsive genes **(B)**, expression of genes associated with lipid metabolism, lipid droplet biology, peroxisomal function, and autophagy **(C)**, expression of endoplasmic reticulum-associated genes **(D)**, apoptosis-related genes **(E)** in hepatocytes from control and ethanol-treated larvae. Dot size represents the percentage of cells expressing a given gene, whereas color intensity indicates the average expression level within each condition.

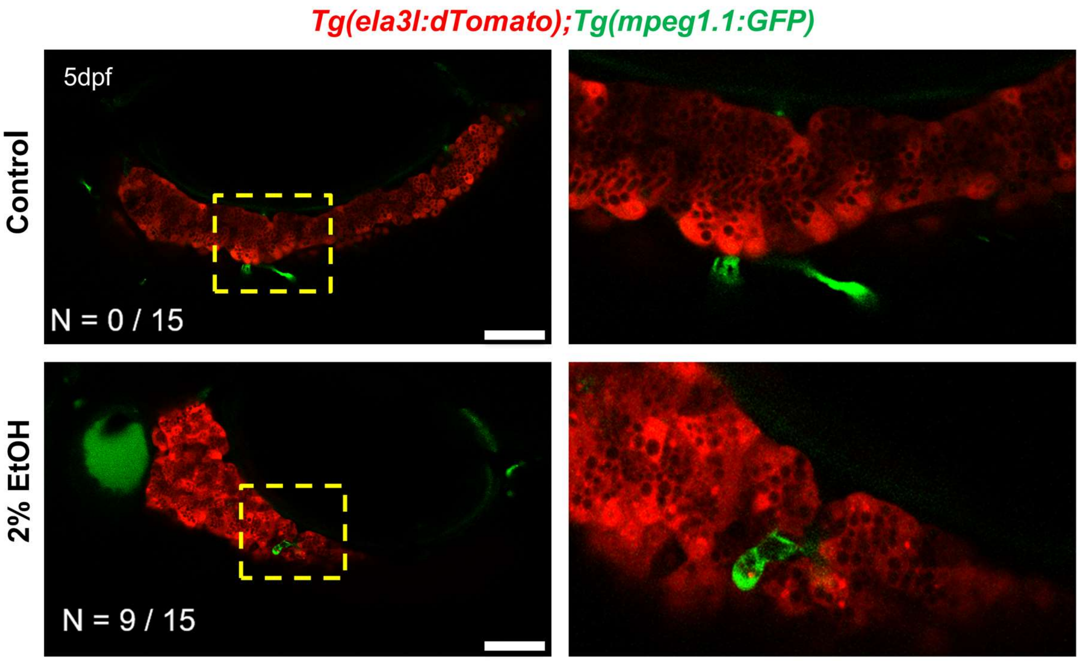


**Supplementary Figure 3: Ethanol exposure induces macrophage infiltration into the pancreas.**

Representative confocal images of 5 dpf *Tg(ela3l:dTomato); Tg(mpeg1.1:GFP)* larvae, in which pancreatic acinar cells are labeled with dTomato (red) and macrophages are labeled with GFP (green). Under control conditions, only a limited number of macrophages were observed in close proximity to the pancreas. Following exposure to 2% ethanol (EtOH), increased accumulation and infiltration of GFP-positive macrophages were detected within the pancreatic tissue. Yellow dashed boxes indicate regions shown at higher magnification in the right panels. Scale bar represents 20μM for all panels.

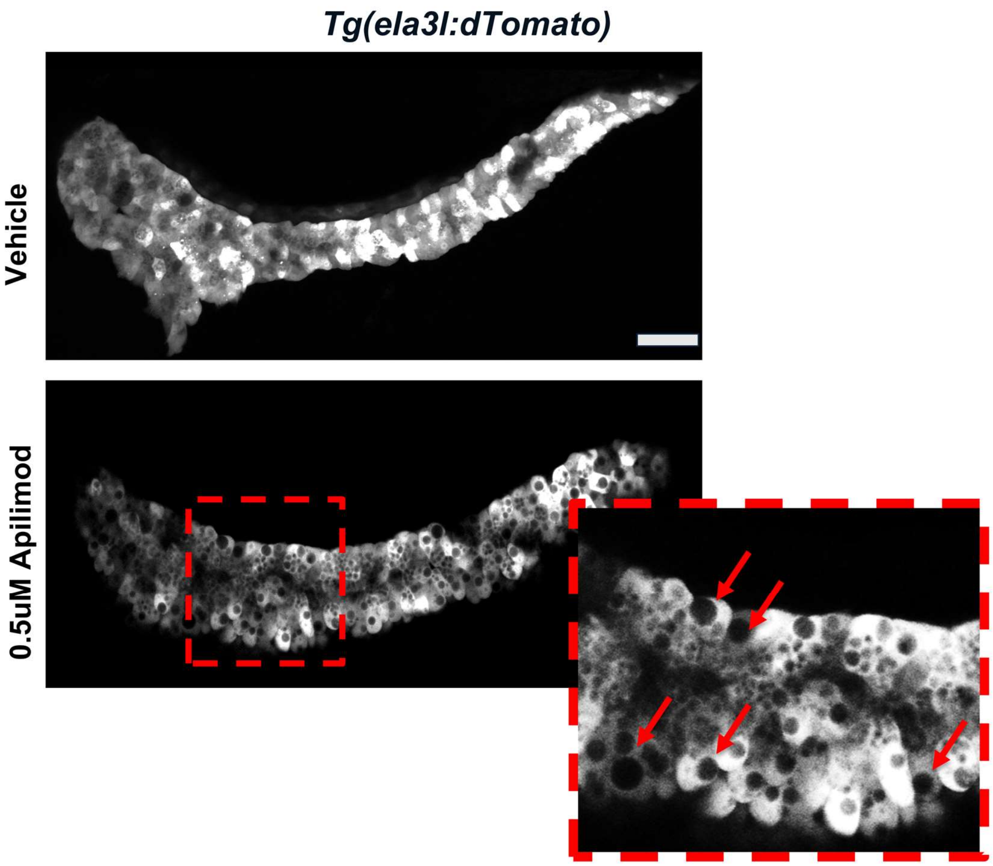


**Supplementary Figure 4: Pharmacological inhibition of PIKfyve induces vacuolar phenotype in pancreatic acinar cells.**

Representative confocal images of pancreas from *Tg(ela3l:dTomato)* larvae treated with vehicle and 0.5 μM Apilimod. While vehicle-treated larvae exhibited normal acinar morphology, Apilimod treatment induced extensive intracellular vacuolation characterized by the accumulation of enlarged vacuolar compartments (red arrows). The boxed region is shown at higher magnification. Scale bars represent 20μM for all panels.
